# PEXMap: A proteogenomic method for exon and isoform level mapping of mass spectrometry derived peptides

**DOI:** 10.64898/2026.04.29.721330

**Authors:** Deepanshi Awasthi, Paras Verma, Shashi Bhushan Pandit

## Abstract

Alternative splicing (AS) expands transcriptome and proteome diversity by differentially combining exons or their splice variants. Although RNA-seq studies have uncovered transcriptomic variability, understanding the corresponding protein-level diversity remains limited. Mass spectrometry-based proteomics provides protein-level insights through MS/MS peptide annotations, which are mostly linked to gene/transcript or UniProt identifiers. However, tracing them to specific isoforms remains challenging due to the lack of exon mapping or inconsistent annotations. We developed PEXMap (**P**eptide**EX**on**Map**per), a k-mer-based proteogenomic framework that systematically maps MS/MS peptides to genes, transcripts, exons, or exon-exon junctions by exact matching of unique 8-mers derived from MS/MS peptides to those in reference databases from exon-resolved isoforms. Comparing PEXMap mappings of human proteome from PeptideAtlas showed annotation concordance with it. Applying PEXMap to liver and pancreas proteomes, we identified tissue-specific isoform expression and, similarly, annotated the cancer proteome. PEXMap reliable mappings could provide insights into role of AS in shaping proteomes across tissues and disease states. Source code is publicly available for download at GitHub: https://github.com/deepanshicbg/PEXMap and supported on Linux.

## Introduction

The phenotypic complexity in higher eukaryotes, especially in humans, emerges from a relatively limited set of protein-coding genes (∼20-22K), which generate diverse proteoforms. These arise from alternative splicing (AS), alternative transcriptional or translational initiation and termination, and post-translational modifications (Aebersold, et al., 2018). Among these, AS is widely recognized as a primary driver of transcriptome or proteome diversity, as has been shown that ∼95% human multi-exon genes produce multiple isoforms (Gonzalez-Porta, et al., 2013; Pan, et al., 2008; Wang, et al., 2008). These transcripts vary in exon composition, coding exon architecture, and regulatory elements, thereby, expanding the functional complexity without increasing gene numbers. The coding exons often encompass functional domains for catalysis/binding, protein-protein interaction, subcellular localization, or post-translational modifications (Kelemen, et al., 2013; Nilsen and Graveley, 2010). Consequently, the inclusion/exclusion of exons or their splice-variants in isoforms can reshape protein architecture expanding gene’s functional repertoire. AS plays a crucial role in various cellular processes such as cell fate determination, development, and adaptive responses (Barash, et al., 2010; Naftelberg, et al., 2015). Aberrant splicing could disrupt functional/regulatory regions in transcripts that results in altered expression/function contributing to diseases (David and Manley, 2010; Sveen, et al., 2016).

RNA-Seq has facilitated systematic characterization of transcriptome diversity, transcript expression, novel splice variant detection, and differential AS patterns across tissues, cellular conditions, and states (Gonzalez-Porta, et al., 2013; Pan, et al., 2008; Wang, et al., 2008). While these provide key insights into splicing-induced transcript variability and its regulation, they are limited in uncovering protein-level consequences. Moreover, comparative transcriptomic and proteomic analyses show moderate mRNA-protein abundance correlations, indicating protein expression is affected by other regulatory processes such as translation efficiency (Liu, et al., 2016; Vogel and Marcotte, 2012). Consequently, transcript detection alone is insufficient to establish isoform expression; rather, protein-level quantification is required to assess splicing-derived proteome diversity. Mass spectrometry (MS) based proteomics provides sequence-resolved peptides and is widely used approach for experimental large-scale protein identification (Nesvizhskii, 2014). Traditional MS methods annotate peptides by matching mass to charge (m/z) ratio spectra to reference spectra, whereas tandem mass spectrometry (MS/MS) determines sequence by fragmenting peptides and is more reliable for protein identification. Thus, MS/MS data is well-suited for analyzing protein expression, specifically for detecting isoform-specific expression. Moreover, integrating proteomics data with genomics and transcriptomics in proteogenomics approaches could improve genome annotation by linking transcript variants to their expressed proteins (Nesvizhskii, 2014; Sheynkman, et al., 2013; Woo, et al., 2014). In proteogenomic approaches, customized protein sequence databases derived from transcriptomic data expand the searchable sequence space and improve detection of novel peptides, alternative splice forms, and mutation-derived variants (Nesvizhskii, 2014). Most of these pipelines primarily map MS/MS peptides to gene, one or multiple transcripts, or protein identifiers (such as UniProt (UniProt, 2025)). Despite these, there are significant challenges in handling ambiguous isoform assignments and heterogeneous annotations, which limit the ability to perform comparative or meta-analyses to detect tissue-/disease-specific isoforms.

To address challenges in mapping MS/MS peptides and standardize annotation for meta-analyses, we developed PEXMap (**P**eptide**EX**on**Map**per), an exon-aware proteogenomic framework to perform systematic annotations of peptides from exons to isoforms/transcripts and genes. Importantly, we utilize unique exon-exon junction (EXj) for isoform-specific detections, though this is constrained by occurrence of unique EXj. Further, we track exon through its nomenclature, which characterize each exon by its relative position in gene, splice variations, and coding sequence (Verma, et al., 2025).

PEXMap involves exact matching of k-mer (8-mer) sequence to map experimental MS/MS peptides to exon-resolved isoform sequences. For this, we constructed distinct overlapping theoretical 8-mer sequences from isoforms annotated in ENACTdb that are compiled into a searchable, indexed reference database (hash table) for efficient lookup, while retaining their associated genes, transcripts/isoforms, exons, and exon-exon junctions. We annotate MS/MS peptide by exact matching of similarly decomposed 8-mers against the indexed reference database followed by propagation of corresponding annotations (gene, transcripts/isoforms, exon) to experimental peptides. Importantly, multi-level mapping is feasible due to trackable coordinates and transcript contexts of exons accessible in ENACT annotations. We assessed the PEXMap annotation on extensive human MS/MS peptide dataset from PeptideAtlas (Desiere, et al., 2006), using its annotation as a reference. We found that PEXMap mappings were highly concordant with those of PeptideAtlas at various levels and could discriminate specific isoforms using exon-exon junctions. Moreover, enabling exon association with peptide could also be used for discriminating among transcripts. Applying PEXMap on liver and pancreas proteome, we could identify tissue-specific expression of isoforms and detected isoforms expressed in cancers by analyzing cancer proteomes.

## 2. Materials and Methods

### 2.1 Data Sources

#### 2.1.1 Human reference genome and transcriptome dataset

We extracted transcripts/isoforms and exon annotations of genes in human genome from ENACT (v.0.5) database (https://www.iscbglab.in/enactdb). This consisted of 19,730 protein-coding genes generating 114,541 transcripts/isoforms (built on NCBI RefSeq GRCh38 reference assembly). Each exon is assigned a unique Exon Unique IDentifier (EUID) characterizing its ordinal position, occurrence, coding status and splice variant features (for details of exon nomenclature, see S1 text (available as supplementary data)). A transcript/isoform is defined as a combination of exons EUIDs that constitute it.

#### 2.1.2 Experimental Mass Spectrometry Data

We obtained experimental human MS/MS peptides from the PeptideAtlas repository (https://peptideatlas.org/builds/human/). We used proteome dataset (Full Build 2022-01) for benchmarking PEXMap. Similarly, we retrieved liver and pancreas proteomics to detect tissue-specific isoforms and pooled peptides from cancer tissue/cells to construct the cancer proteome.

We filtered MS/MS peptides to remove identical sequences, sequences of length <8, and homopolymer repeats (Shannon entropy=0). Further, peptides associated with multiple genes were considered ambiguous and excluded from the analysis. The filtering step reduced the initial human proteome from 2,883,406 peptides to 1,739,961. Similarly, filtered MS/MS peptides in the liver and pancreas proteome were 184,342 and 75,517, respectively. The pooled cancer proteome comprised of 1,152,437 processed peptides.

#### 2.2 Overview of PEXMap

The k-mer based approaches are efficient at matching protein or DNA sequences, especially for large-scale sequence comparisons (Moeckel, et al., 2024). Usually, these involve segmenting sequences into k-mers, which are then compared against precomputed indexes to filter matches at low computational cost. Some implementations of k-mer based methods include PEPMatch (Marrama, et al., 2023), BLAT (Kent, 2002), DIAMOND (Buchfink, et al., 2015), and MMSeq2 (Steinegger and Soding, 2017).

Considering k-mer efficiency, we developed PEXMap to associate MS/MS peptides with genes, isoforms/exons by identifying shared k-mer sequences. We choose *k=8* because it efficiently resolves and captures exon boundaries, provides peptide coverage across transcripts for specificity, and reduces spurious string matching. Below, we briefly describe the construction of an 8-mer reference-indexed database, querying of 8-mers extracted from peptides, and their multi-level mapping. An overview of PEXMap is illustrated in Fig. 1.

**Figure 1:**
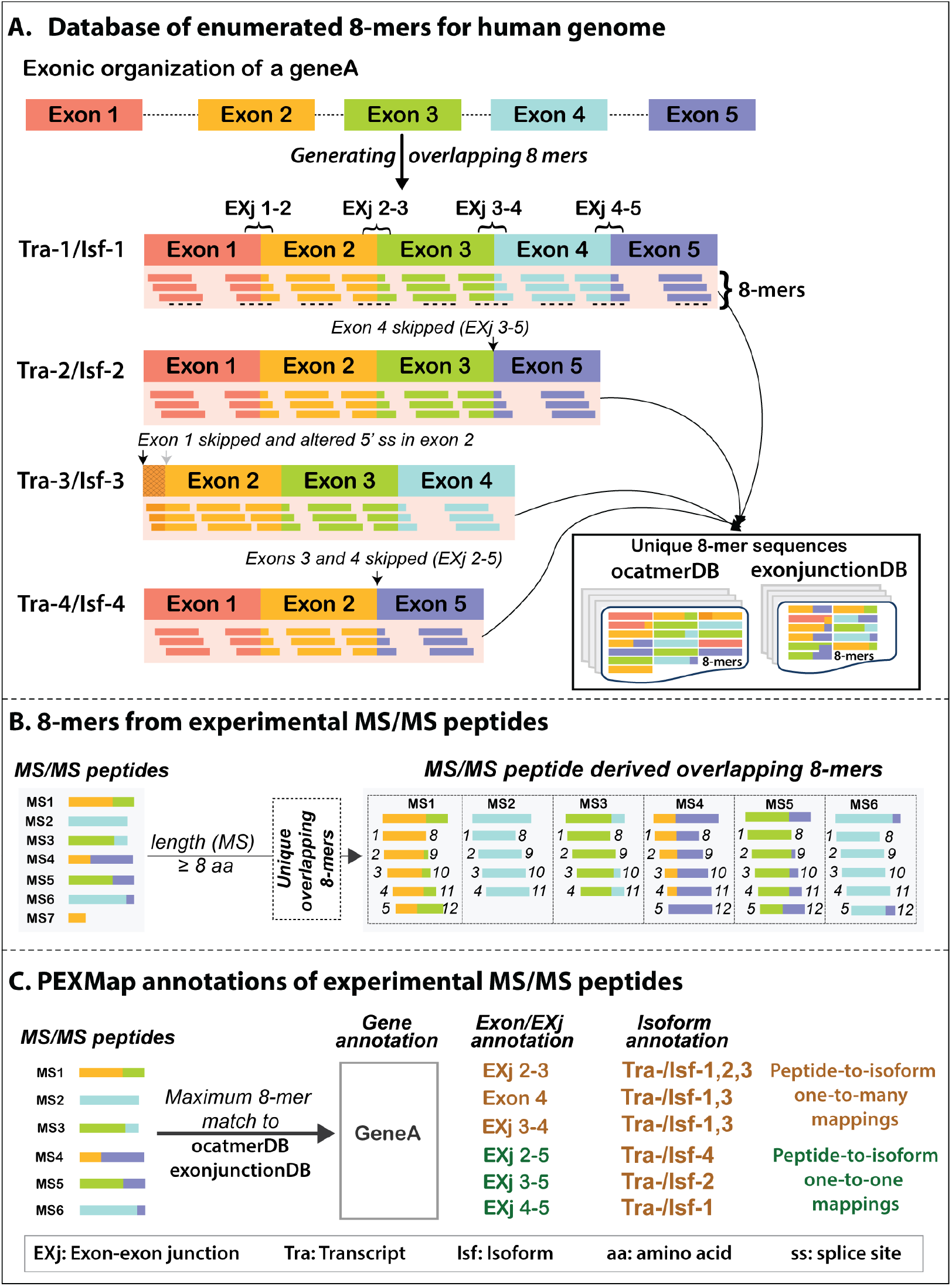
Overview of PEXMap includes construction of k-mers. Schematic illustrations showing generation of overlapping 8-mer in: (A) to construct reference databases from isoforms (octamerDB) and similarly for exon-exon junction (EXj) across isoforms (exonjunctionDB) and (B) for experimental MS/MS peptides from PeptideAtlas. (C) Workflow illustrating exact matching of experimental peptide derived 8-mers to reference database followed by multi-level annotation of peptides with gene, isoforms(s), exon(s) and EXj(s).

#### 2.2.1 Construction of 8-mer sequence library for reference database and MS/MS peptides

As efficient searching of 8-mers requires an indexed reference database, we constructed a theoretical 8-mer overlapping peptide sequence database (octamerDB) from human isoforms (ENACTdb). Briefly, each isoform is fragmented into overlapping 8-mer sequences using a one-residue offset. The resulting unique 8-mer sequences were collated for a gene and mapped to their corresponding gene/isoform(s). Some theoretical peptides may map to multiple isoforms due to shared sequences. Considering exon-resolved isoform sequence derived from ENACT, each 8-mer was further linked to either an exon or exon-exon junction (EXj). The latter corresponds to peptides spanning two adjacent exons in an isoform that we referred to as ‘EXj-peptides’. Unique EXjs enable discriminating among isoforms, as it defines a specific exon-exon combination. Because exons are identified by their unique identifier (EUID), we can track multi-mapping to resolve splice-site variations.

Next, we collated unique 8-mer sequences from all human genes, traced their provenance to gene(s)/isoform(s)/exon(s), and indexed them in octamerDB as a hash lookup data structure. Similarly, we created a dedicated 8-mer indexed database (exonjunctionDB) of EXjs derived from isoform(s). We had a total of 1,363,811 8-mers after combining unique exon-exon junction 8-mers across the human genes. Of these, 53,827 mapped to more than one gene.

To construct 8-mers for experimental peptides, we followed the same steps as for database construction, retaining unique peptides of length ≥ 8 and without repeating amino acids (Fig. 1B). Each 8-mer was retained with its corresponding identifier for tracking mappings.

#### 2.2.2 Method to annotate MS/MS peptides

To assign experimental peptides at each level, we performed exact sequence matches between 8-mers derived from MS/MS peptides and those in the reference database. As 8-mers were stored in a hash table, it essentially served as a lookup of 8-mers from MS/MS peptides in both indexed databases (octamerDB and exonjunctionDB). Subsequently, we assigned MS/MS peptides to a gene/isoform/exon/EXjs based on the maximum number of matching 8-mers between experimental peptides and the reference database. We conducted the above annotation in two stages. First, we mapped experimental peptides to gene(s), and then assigned them to their corresponding isoform(s), exons(s), and/or EXj(s) (Fig. 1C). At each stage of peptide assignment, we used the maximum-matched 8-mer criterion as mentioned above. However, when multiple genes/isoforms/exons share an equal number of maximal-matching 8-mers, they were assigned accordingly and are referred to as multi-mapping. The latter was common for isoform assignment because sequences encompassing constitutive exons or splice-site variation yield mostly the same 8-mers. Finally, MS/MS peptides were assigned with gene, transcript/isoform identifiers, or exon identifier (EUID) to preserve the multi-level nature of the mapped annotations. It is important to note that we have used exonjunctionDB for assigning MS/MS peptides to EXj(s), enabling peptide annotation to respective isoform(s).

### 2.3 Benchmarking of PEXMap

Gene or transcript annotations of experimental peptides from PeptideAtlas served as a reference for benchmarking PEXMap, and we assessed the concordant annotations between PEXMap and PeptideAtlas. As PeptideAtlas has heterogeneous annotations for gene/transcript identifiers, we carefully cross-referenced them with RefSeq identifiers. Of 15,855 unique genes in PeptideAtlas, we cross-referenced 15,771 and similarly referenced 97,443 transcripts of 112,312.

To assess concordance at the gene level, we selected experimental peptides uniquely assigned to a single gene identifier in PeptideAtlas. We considered a PEXMap mapping accurate if it unambiguously mapped to the same gene identifier in PeptideAtlas. For transcript-level benchmarking, we considered a mapping accurate when all transcript identifiers assigned by PEXMap matched those in PeptideAtlas. Peptides uniquely assigned to single transcripts were analyzed separately to evaluate the ability of the method to discriminate among isoforms. Following the same approach, we independently evaluated gene- and transcript-level assignment performance of PEXMap using exonjunctionDB, which assesses annotation based on EXjs alone. We defined gene/transcript-level accuracy as the proportion of MS/MS peptides in PEXMap that had identical annotations to those reported in PeptideAtlas.

As PeptideAtlas lacks exon-level annotations, we performed an indirect evaluation by first assessing the consistently mapped MS/MS peptides at the gene level and subsequently determining whether they were uniquely assigned to a single exon. The proportion of peptides that supported unique exon-level associations was calculated to assess the resolution of exon-level mapping.

### 2.4 PEXMap for tissue-specific isoform detection and in cancer proteomics

To investigate liver-and pancreas-specific transcripts, MS/MS peptides corresponding to these tissues were obtained from PeptideAtlas and mapped using the PEXMap pipeline. We mapped experimental peptides from each tissue at the gene, transcript, exon, and EXj levels. Further, genes common to both tissues are presumed to be expressed in both and processed further to detect tissue-specific isoforms, such as those mapped to MS/MS peptides in either tissue. Similarly, we used EXjs to detect tissue-specific isoforms.

To identify cancer-associated isoforms, we annotated unique pooled MS/MS peptides from cancer tissue/cell samples in PeptideAtlas, after removing those linked to multiple genes or transcripts. The filtered peptides were assigned to transcripts, exons, and EXj using the PEXMap pipeline. We compared gene level association performance with that reported in PeptideAtlas. Moreover, uniquely assigned exons to MS/MS peptides were examined to identify potential cancer-associated exon usage patterns.

### 2.5 PEXMap implementation

We implemented PEXMap using custom Python scripts. The workflow consists of a preprocessing step to construct 8-mer reference databases, which is performed only once for a genome. The overlapping 8-mers are stored in a Python dictionary for efficient exact matching, and the corresponding gene/isoform/exon annotations are maintained for easy extraction. As k-mer-based approaches are computationally time-efficient, we measured the time required for database construction and searching 8-mers (∼10^7^), which were ∼4 minutes and ∼2 minutes, respectively. Together, these results demonstrate that PEXMap pipeline is computationally efficient and well-suited for larger-scale analysis. The source code and usage instructions are publicly available on GitHub: https://github.com/deepanshicbg/PEXMap

## 3. Results

We constructed (see methods) indexed 8-mer reference databases: octamerDB, derived from 114,541 human isoforms, and a complementary library of 8-mers exonjunctionDB, representing from sequences spanning only adjacent exons from human isoforms. Each entity in the reference database is associated with its corresponding gene, transcript/isoform, and exon identifiers. Below, we discuss benchmarking of MS/MS peptides using PEXMap and its application in detecting tissue-specific isoforms and also in cancer proteomes. The statistics of MS/MS peptide mapping in PeptideAtlas and benchmarking results are summarized in supplementary tables 1, 2 and 3.

### 3.1 Benchmarking of peptide mapping performance

#### 3.1.1 Gene and transcript/isoform level mapping of peptides

We first evaluated PEXMap gene level assignment concordance for 1,739,961 MS/MS peptides uniquely assigned to a single gene in PeptideAtlas (see methods). Using octamerDB, PEXMap accurately mapped 99.4 % of total mapped peptides to their respective genes (Fig. 2A). The remaining peptides either showed multimapping or remained unmapped due to the absence of genes in our dataset. Searching against EXj’s (exonjunctionDB), 95% of total mapped peptides were consistent with gene associations of PeptideAtlas and those spanned ∼78% of genes. Although EXj peptides constitute a smaller fraction of total peptides, the mapping accuracy remained high, demonstrating the reliability of junction-aware annotation. Overall, these results show that PEXMap achieves robust gene-level mapping across large-scale proteomic datasets.

**Figure 2:**
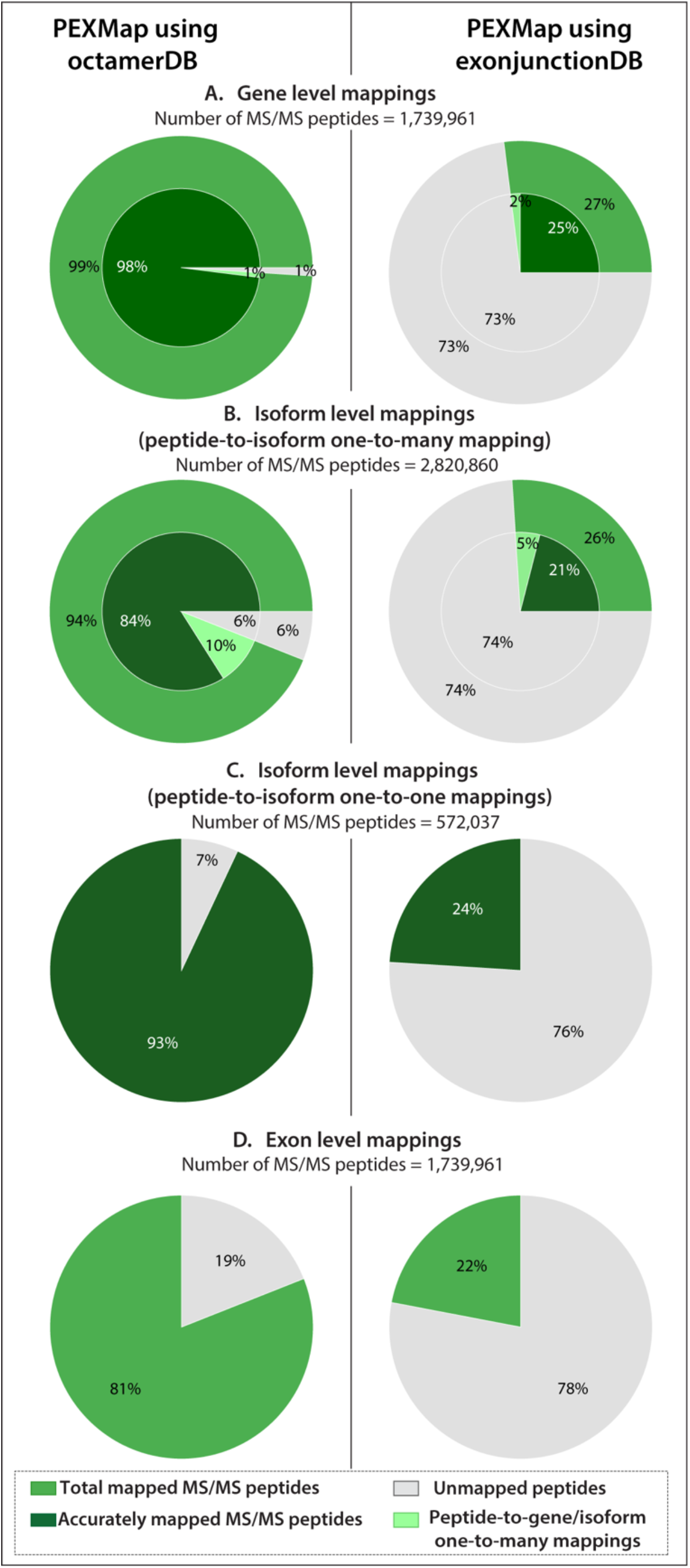
Benchmarking performance of PEXMap on MS/MS peptides from PeptideAtlas. Pie chart summarizing results of experimental peptides mapped by PEXMap using octamerDB (left panel) and EXj-derived exonjunctionDB (right panel). Panels A–D show the percentage distribution at following levels: (A) Gene level, (B) Protein (isoform) level for all peptides, (C) Protein (isoform)-level for uniquely assigned peptides, and (D) Exon-level (specific exons or EXjs). Mapped peptides are shown in green, accurately mapped in dark green, and unmapped in grey. Total number of MS/MS peptides for each mapping level is included appropriately in the figure.

Next, we assessed transcript-level annotations on 2,820,860 MS/MS peptides mapped to transcripts in PeptideAtlas. Of these, 572,037 peptides were associated with a single transcript i.e. peptide-to-isoform with one-to-one mappings, while the remaining were multi-mapped i.e. peptide-to-isoform with one-to-many mappings. Using PEXMap with octamerDB, 84.1% of experimental peptides mapping to transcript annotations were consistent with those in PeptideAtlas (Fig. 2B). It is important to note that, for peptide-to-isoform one-to-many mappings, we considered an assignment by PEXMap to be correct when it matches all listed transcripts for a peptide. Most PEXMap mappings were labelled as inconsistent because they mapped to a subset of listed transcripts. By restricting analysis to peptide-to-isoform one-to-one mappings, we accurately mapped ∼93% of them (Fig. 2C), exhibiting that PEXMap assignment of MS/MS peptides to genes/transcripts were highly concordant to PeptideAtlas.

Considering EXj peptides with PEXMap, ∼27% of MS/MS peptides could be mapped to transcripts; of these, ∼79% were consistent with annotations in PeptideAtlas. We expect a relatively low mapping fraction, as peptides spanning exon-exon junctions are less abundant and are challenging to capture experimentally. Despite junction peptides representing a smaller fraction of total mappings, they provided enhanced reliability for transcript-level discrimination. Focusing on peptides singly mapped to transcripts, ∼25% of peptide assignments were consistent as annotated in PeptideAtlas (Fig. 2C). Notably, the assignment based on it would have higher confidence, as it uses unique exon-exon junctions to characterize isoforms.

These results demonstrate that PEXMap enables reliable isoform-level annotation and that junction-derived peptides improve transcript-specific detection, thereby enabling higher transcript-level precision by resolving isoform-level ambiguities.

#### 3.1.2 Exon-Level Mapping of MS Peptides

Unlike PeptideAtlas, our method also provides exon-level annotation for experimental peptides (Fig. 2D). We mapped 17,12,990 MS/MS peptides to exons. Of these, we assigned 81.8% of peptides to single exons using octamerDB, and using exonjunctionDB, we assigned ∼22% to EXjs or their corresponding genes. Such exon-level resolution provides direct evidence of translated exon usage and demonstrates the significance of exon-aware proteogenomic analysis. To our knowledge, such comprehensive exon-resolved peptide mapping is not currently available in public proteomic repositories.

### 3.2 Application of PEXMap for identifying tissue-specific transcripts

Having demonstrated PEXMap’s ability to annotate MS/MS peptides with exons, transcripts, and isoforms, we applied it to identify tissue-specific isoforms. As a representative analysis, we examined isoform-specific expression in liver and pancreas proteomic data, as these tissues share a substantial fraction of commonly expressed genes while exhibiting distinct physiological functions and regulatory profiles (Ber, et al., 2003; Jia, et al., 2026; Zaret and Grompe, 2008).

We performed multi-level mapping using PEXMap of experimental peptides (see methods) from liver/pancreas (Supplementary Table S2, available as supplementary data). At the gene level, we identified 8,064 and 6,189 genes in the liver and pancreas, respectively. Focusing on identifying tissue-specific isoforms, we considered only genes (5,774) expressed in both tissues. Among these genes, ∼81% exhibited MS/MS peptides (4,698) mapped to identical transcripts in liver and pancreas (octamerDB-based searches). Such shared peptides from transcripts could arise from constitutive exons or exon-exon junctions shared across multiple isoforms. From the remaining mappings (∼19%), we extracted genes with at least one unique transcript identified in either tissue. To ensure robustness, we quantified the number of MS/MS peptides mapped to these unique isoforms and checked for distinguishing exons or EXj between isoforms. Based on these, we identified 1,076 genes with differential transcript usage between both tissues. Of these genes, liver showed 273 genes with unique transcripts, comprising 1,129 isoforms supported by 1,175 unique peptides (∼4 peptides per gene). On contrary, pancreas showed 58 genes with 181 isoforms supported by 80 unique peptides (∼1 peptide per gene). Further, many isoforms were supported by EXj peptide mapping and transcript-specific unique peptides, indicating tissue-specific isoform expression. Notably, the liver showed greater peptide support and a greater number of uniquely supported transcripts than the pancreas, reflecting higher proteomic coverage. These findings demonstrate that even when tissues express the same genes, they differed at the level of isoform expression. This is consistent with previous transcriptomic and proteogenomic studies showing that AS is tissue-specific and a major contributor to proteome diversity (Consortium, 2020; Pan et al., 2008; Wang et al., 2008). Below, we discuss several examples of tissue-specific transcript expression supported by the presence of EXj peptides.

#### MYL6

*MYL6* (NCBI ID: 4637) encodes myosin light chain, which generates distinct isoforms through AS, and differential exon usage results in functionally different variants across cell types (Lenz, et al., 1989). Myosin is a highly conserved contractile protein present in muscle and non-muscle cells. It exists as part of a diverse multigene family, with isoform transitions regulated during development and disease (Bandman, 1985). Exon-6 inclusion in MYL6 shows strong cell-type specificity, with higher inclusion in tumor epithelial cells and reduced inclusion in immune cells (Fu, et al., 2025). We found exon-6 containing isoform (NP_066299.2) in liver supported by exonic and EXj peptides (n=32) from EXjs spanning exon-5 and exon-6 (Fig. 3A). In contrast, exon-6 is skipped in pancreas isoform (NP_524147.2). Accordingly, we found peptides (n=18) spanning exon-5 and exon-7 (Fig. 3A). Notably, both isoforms encode proteins of equal length, as inclusion of exon-6 introduces a premature termination and excludes exon-7 from the coding region. This difference in *MYL6* tissue-specific isoform expression likely contributes to its observed cell-type-specific features and may be relevant to disease-associated splicing variation.

**Figure 3:**
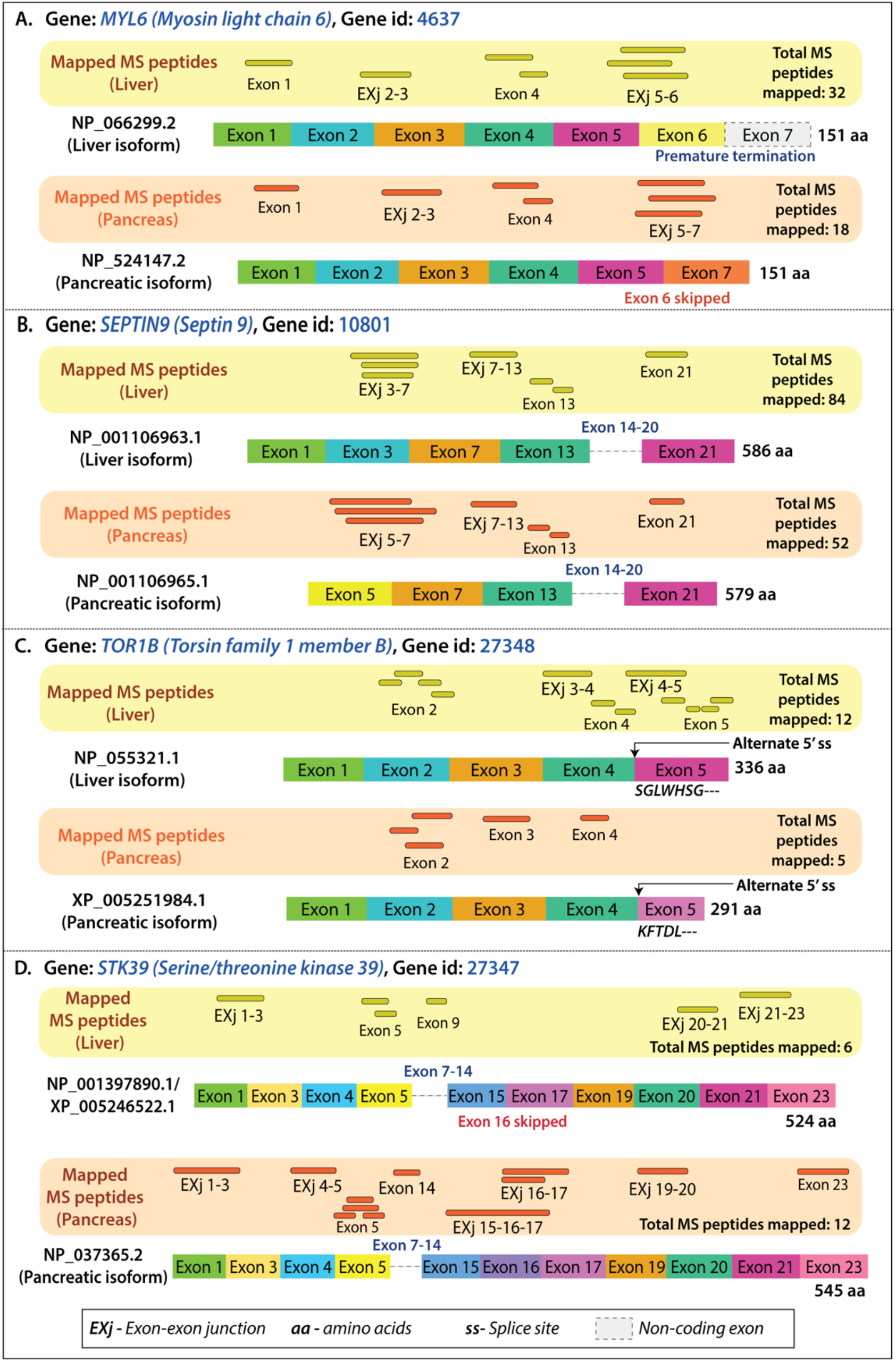
Representative genes showing tissue-specific isoform expression. Each panel shows MS/MS peptides mapping at the isoform level alongside their respective exon organization for representative genes. (A) *MYL6:* isoform (NP_066299.2) expressed in liver includes exon-6, which is skipped in pancreas-specific isoform (NP_524147.2). (B) *SEPTIN9* isoforms expressed in liver (NP_001106963.1) and pancreas (NP_001106965.1) differ by alternative internal exon usage. (C) *TOR1B* isoforms (liver: NP_055321.1, pancreas: XP_005251984.1) vary due to alternative 5′ splice site selection. (D) *STK39* liver isoform (NP_001397890.1) varies from pancreatic isoform (NP_037365.2) by alternative exon usage, including exon-16 skipping. In each panel, annotated isoforms are shown as combinations of exons represented by rectangular exon boxes, with splice-site variations indicated by altered exon boundaries. Matched MS/MS peptides are displayed as thick lines above the corresponding isoform and numbers indicate the total mapped peptides supporting each isoform. Yellow and orange boxes represent liver- and pancreas-derived peptides, respectively. EXj denotes exon-exon junction peptides, “ss” indicates splice site, and dashed boxes indicate non-coding exons.

#### SEPTIN9

*SEPTIN9* (NCBI ID: 10801) gene encodes a cytoskeletal GTP-binding protein involved in membrane remodeling and cell division (Mostowy and Cossart, 2012). Liver showed expression of isoform (NP_001106963.1) as supported by multiple MS/MS peptides to EXjs (e.g., EXj 3–7 and EXj 7–13) and other (n=84) mapped peptides (Fig. 3B). In contrast, a shorter isoform (NP_001106965.1) is expressed in pancreas supported by pattern of EXj peptides (e.g., EXj 5–7 and EXj 7–13) and other (n=52) exonic peptides (Fig. 3B). Given that septin-mediated pathways are implicated in liver disease and tumor progression (Fan, et al., 2021), these findings highlight significance of PEXMap in detecting isoform-specific expression or enrichment in disease contexts by providing resolution beyond conventional gene-level analysis.

Genes *SCAMP3* (NCBI ID: 10067) and *ACOX1* (NCBI ID: 51) showed similar tissue-specific isoform expression supported by isoform-specific exons, and EXj peptides are discussed in S2 text (available as supplementary data).

In the above discussed genes (Figs. 3A-B), peptides were mapped to unique EXj in either tissue supporting the isoform-specific expression; however, it is not always feasible to identify such EXj-spanning peptides for isoform-specific gene expressions. Below, we discuss such genes, with evidence at exonic level or unique EXj evidence in either tissue.

*TOR1B* gene (NCBI ID: 27348), a member of the torsin protein family (Ozelius, et al., 1997), plays an important role in maintaining cellular homeostasis and mediating cellular response to endoplasmic reticulum (ER) stress (Zhang, et al., 2024). It belongs to the AAA+ protein family (InterPro ID: IPR003959), which is involved in diverse cellular processes, including protein folding, trafficking, and degradation within ER (Rose, et al., 2015). MS/MS peptides annotated in liver dataset (Fig. 3C) supported the full-length isoform (NP_055321.1) with clear EXj evidence. In contrast, pancreas samples showed limited peptide coverage for a shorter isoform (XP_005251984.1), with no peptides mapped to EXjs.

We observed a similar pattern for *STK39* gene, also known as *SPAK* and involved in kinase signaling pathways (Vitari, et al., 2005), where peptides corresponding to exon-16 and its flanking junctions were detected only in pancreas (Fig. 3D), supporting the isoform (NP_037365.2), while liver does not have MS/MS peptides mapped to this junction limiting precise isoform assignment despite peptide detection.

Together, these examples highlight PEXMap’s ability to move beyond gene-level detection and enable isoform-resolved interpretation of proteomic data, where junction (EXj) peptide evidence provides the most reliable support for confident transcript identification, offering a more precise view of tissue-specific expression in liver and pancreas.

### 3.3 Isoform mapping in cancer proteome

We evaluated PEXMap on a pooled cancer peptide dataset, comprising 2,104,053 unique experimental peptides derived from cancer tissues and cell lines. After removing peptides with ambiguous annotations, we associated 1,152,437 peptides with 14,549 genes and 326,423 peptides with 8,417 transcripts based on PeptideAtlas records (Supplementary Table S3, available as supplementary data). Using PEXMap, we found high concordance with PeptideAtlas with 97% of peptides mapped to 14,296 genes and 89% of peptides to 7,489 transcripts. Notably, we could also link 81% of the peptides to specific exons, thereby revealing peptides that support cancer-associated isoforms. Below, we discuss two well-known cancer-associated genes, *EGFR and FLNA*, as a case study in which literature-supported isoform changes link to functional variation.

#### EGFR

*EGFR* (NCBI ID: 1956) gene encodes a key oncogenic receptor that, upon ligand binding, activates its intracellular tyrosine kinase domain (toward the protein C-terminus) to drive cell proliferation and survival pathways (Voldborg, et al., 1997). Pooled cancer dataset showed MS/MS peptides mapping to EXj 17-19 (Fig. 4A) with no observation of peptides mapped to downstream exons, indicating expression of a truncated isoform (NP_958441.1), which lacks the catalytic domain. This observation is consistent with well-established *EGFR* alterations in cancer, particularly exon-19 variants that affect kinase activity and therapeutic response (Lynch, et al., 2004; Paez, et al., 2004).

**Figure 4:**
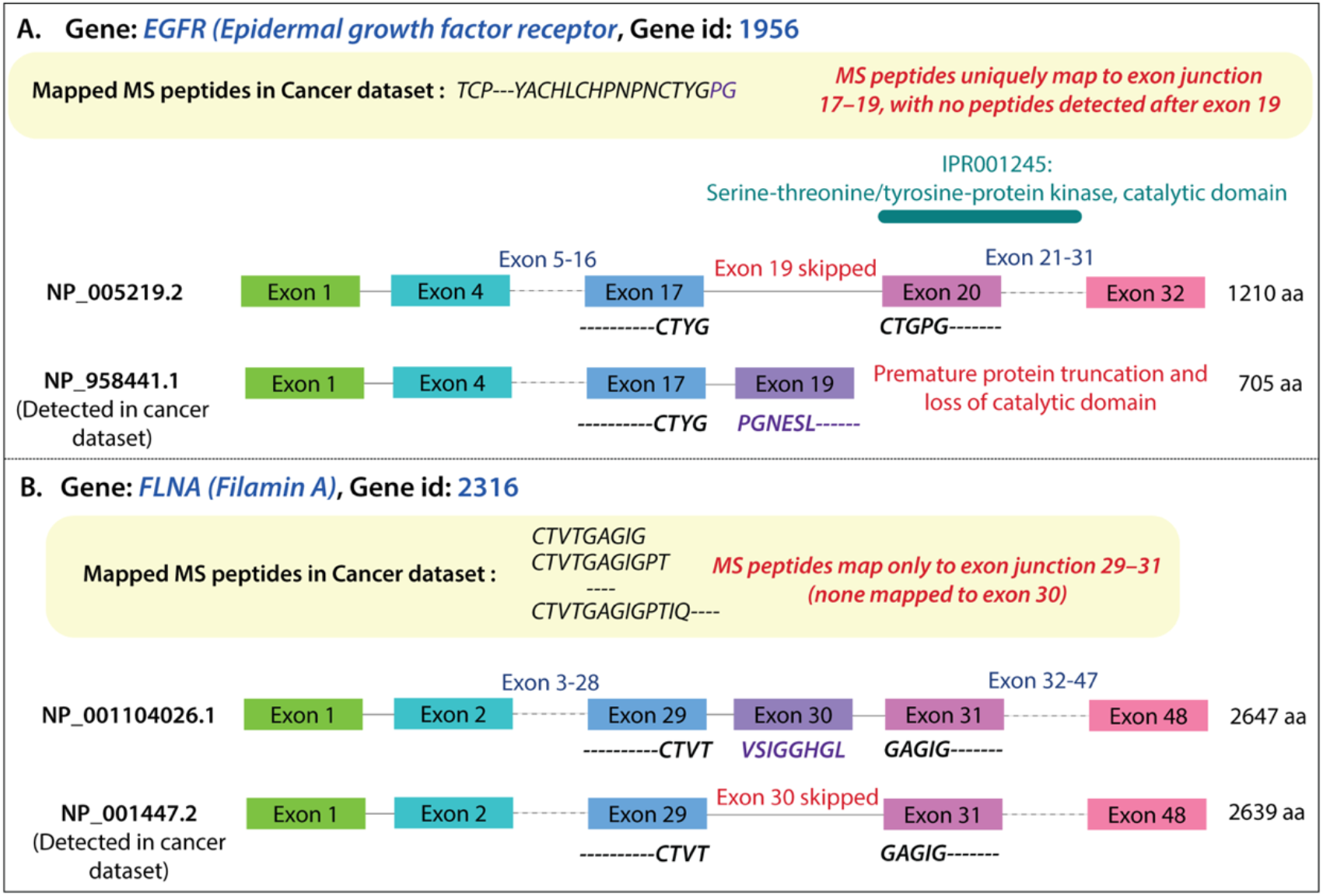
Representative genes showing isoform-specific expression in cancer proteome. (A) The *EGFR* isoform (NP_958441.1) in the cancer proteome has MS/MS peptides mapped to exon junctions (EXj 17–19) but lacks downstream peptide evidence, indicating a truncated isoform missing the C-terminal kinase domain compared to the full-length isoform (NP_005219.2). (B) The *FLNA* isoform (NP_001447.2) contains peptides spanning exon junctions (EXj 29–31) but lacks support for exon-30, indicating an exon-30-skipped isoform compared to the canonical isoform (NP_001104026.1). Peptide support and alternative splicing are highlighted in red, following the schematic format in Fig. 3.

#### FLNA

The *FLNA* (NCBI ID: 2316) gene encodes an actin-binding scaffold protein that regulates cytoskeletal organization, cell adhesion, migration, and multiple signaling pathways implicated in tumor progression (Zhou, et al., 2021). Consistently, we observed peptides spanning the EXj 29–31 with no peptides mapping to exon-30 (Fig. 4B), directly supporting an exon-30 skipped isoform (NP_001447.2) in cancer. Notably, exon-30 encodes part of a filamin repeat domain critical for cytoskeletal remodeling, and its AS has been associated with tumor progression and altered cell motility (Lu, et al., 2015; Savoy and Ghosh, 2013).

Another gene, *CASP2* (NCBI ID: 835), demonstrating isoform-specific peptide mapping in cancer, shown in S3 text (available as supplementary data).

The above study demonstrates our approach’s ability to resolve functionally relevant splice variants. Together, these examples illustrate how PEXMap leverages experimental MS peptides to move beyond gene-level detection and directly identify the isoforms present in cancer dataset through exon-level evidence.

## 4. Discussion

AS extensively diversifies transcriptome and proteome, yet confirming isoform-specific expression at the protein level remains challenging. We developed PEXMap, a k-mer based method to systematically annotate MS/MS peptides obtained from mass spectrometry-based proteomics. We mapped peptides at multiple levels, from genes and transcripts to exons and exon–exon junctions. Benchmarking PEXMap on PeptideAtlas annotations demonstrated high concordance and coverage with EXj mapping as an important feature to discriminate among isoforms. We applied PEXMap to liver and pancreas proteomes, showing tissue-specific isoform expression, and similarly provided isoform-level annotations in cancer proteomes. PEXMap can be applied to annotate other proteomes; however, this depends on the level of isoform annotation and abundance of MS/MS peptides. Overall, PEXMap provides a scalable framework for analyzing how alternative splicing shapes proteomes across tissues and disease states.

## Acknowledgements

We acknowledge IISER Mohali for initial financial support. Ms. Deepanshi Awasthi acknowledges Prime Minister’s Research Fellowship [PMRF ID: 0601560] for financial support during Ph.D.

## Author Contributions

D.A. (Methodology [lead], Software [lead], Formal analysis [lead], Investigation [lead], Writing—original draft [lead], Writing—review & editing [supporting]), P.V. (Resources [lead], Writing—review & editing [supporting]), and S.B.P. (Conceptualization [lead], Supervision [lead], Methodology [Supporting], Investigation [supporting], Writing—review & editing [lead]).

## Supplementary Material

Supplementary material is available as supplementary data.

## Data Availability

The PEXMap source code and usage instructions are publicly available at GitHub: https://github.com/deepanshicbg/PEXMap

The datasets analysed in this study were obtained from publicly accessible repositories, including PeptideAtlas. Processed datasets and intermediate files generated during the current study are available from the corresponding author upon request.

## Supplementary data

S1. **Details of exon nomenclature using ENACT framework**

S2. **Tissue-specific isoforms of *SCAMP3* and *ACOX1* supported by peptide mapping**.

Supplementary Figure S1. **Liver- and pancreas-specific isoforms in representative genes**.

S3. ***CASP2* isoform supported by exon-level peptide evidence in cancer**

Supplementary Figure S2. ***CASP2* isoform in cancer proteome**.

Supplementary Table S1. **Detailed statistics of MS/MS peptides multi-level mapping to gene/transcript/exon/exon-exon junction for each of 8-mer database**.

Supplementary Table S2. **Detailed statistics of MS/MS peptides matched to liver and pancreas proteome**.

Supplementary Table S3. **Detailed statistics of MS/MS peptides matched to cancer proteome**.

## References

Aebersold, R., et al. How many human proteoforms are there? Nat Chem Biol 2018;14(3):206– 214.

Bandman, E. Myosin isoenzyme transitions in muscle development, maturation, and disease. Int Rev Cytol 1985;97:97–131.

Barash, Y., et al. Deciphering the splicing code. Nature 2010;465(7294):53–59.

Ber, I., et al. Functional, persistent, and extended liver to pancreas transdifferentiation. J Biol Chem 2003;278(34):31950–31957.

Buchfink, B., Xie, C. and Huson, D.H. Fast and sensitive protein alignment using DIAMOND. Nat Methods 2015;12(1):59–60.

David, C.J. and Manley, J.L. Alternative pre-mRNA splicing regulation in cancer: pathways and programs unhinged. Genes Dev 2010;24(21):2343–2364.

Desiere, F., et al. The PeptideAtlas project. Nucleic Acids Res 2006;34(Database issue):D655– 658.

Fan, Y., et al. SEPT6 drives hepatocellular carcinoma cell proliferation, migration and invasion via the Hippo/YAP signaling pathway. Int J Oncol 2021;58(6).

Fu, Y., et al. Single cell and spatial alternative splicing analysis with Nanopore long read sequencing. Nat Commun 2025;16(1):6654.

Gonzalez-Porta, M., et al. Transcriptome analysis of human tissues and cell lines reveals one dominant transcript per gene. Genome Biol 2013;14(7):R70.

Jia, Y., et al. Liver-pancreas communication in disease and drug development. Acta Pharm Sin B 2026;16(3):1250–1271.

Kelemen, O., et al. Function of alternative splicing. Gene 2013;514(1):1–30.

Kent, W.J. BLAT--the BLAST-like alignment tool. Genome Res 2002;12(4):656–664.

Lenz, S., et al. The alkali light chains of human smooth and nonmuscle myosins are encoded by a single gene. Tissue-specific expression by alternative splicing pathways. J Biol Chem 1989;264(15):9009–9015.

Liu, Y., Beyer, A. and Aebersold, R. On the Dependency of Cellular Protein Levels on mRNA Abundance. Cell 2016;165(3):535–550.

Lu, Z.X., et al. Transcriptome-wide landscape of pre-mRNA alternative splicing associated with metastatic colonization. Mol Cancer Res 2015;13(2):305–318.

Lynch, T.J., et al. Activating mutations in the epidermal growth factor receptor underlying responsiveness of non-small-cell lung cancer to gefitinib. N Engl J Med 2004;350(21):2129– 2139.

Marrama, D., et al. PEPMatch: a tool to identify short peptide sequence matches in large sets of proteins. BMC Bioinformatics 2023;24(1):485.

Moeckel, C., et al. A survey of k-mer methods and applications in bioinformatics. Comput Struct Biotechnol J 2024;23:2289–2303.

Mostowy, S. and Cossart, P. Septins: the fourth component of the cytoskeleton. Nat Rev Mol Cell Biol 2012;13(3):183–194.

Naftelberg, S., et al. Regulation of alternative splicing through coupling with transcription and chromatin structure. Annu Rev Biochem 2015;84:165–198.

Nesvizhskii, A.I. Proteogenomics: concepts, applications and computational strategies. Nat Methods 2014;11(11):1114–1125.

Nilsen, T.W. and Graveley, B.R. Expansion of the eukaryotic proteome by alternative splicing. Nature 2010;463(7280):457–463.

Ozelius, L.J., et al. The early-onset torsion dystonia gene (DYT1) encodes an ATP-binding protein. Nat Genet 1997;17(1):40–48.

Paez, J.G., et al. EGFR mutations in lung cancer: correlation with clinical response to gefitinib therapy. Science 2004;304(5676):1497–1500.

Pan, Q., et al. Deep surveying of alternative splicing complexity in the human transcriptome by high-throughput sequencing. Nat Genet 2008;40(12):1413–1415.

Rose, A.E., Brown, R.S. and Schlieker, C. Torsins: not your typical AAA+ ATPases. Crit Rev Biochem Mol Biol 2015;50(6):532–549.

Savoy, R.M. and Ghosh, P.M. The dual role of filamin A in cancer: can’t live with (too much of) it, can’t live without it. Endocr Relat Cancer 2013;20(6):R341–356.

Sheynkman, G.M., et al. Discovery and mass spectrometric analysis of novel splice-junction peptides using RNA-Seq. Mol Cell Proteomics 2013;12(8):2341–2353.

Steinegger, M. and Soding, J. MMseqs2 enables sensitive protein sequence searching for the analysis of massive data sets. Nat Biotechnol 2017;35(11):1026–1028.

Sveen, A., et al. Aberrant RNA splicing in cancer; expression changes and driver mutations of splicing factor genes. Oncogene 2016;35(19):2413–2427.

UniProt, C. UniProt: the Universal Protein Knowledgebase in 2025. Nucleic Acids Res 2025;53(D1):D609–D617.

Verma, P., et al. Exon Nomenclature And Classification of Transcripts (ENACT) provides a systematic framework to annotate exon attributes. Genome Res 2025;35(6):1440–1455.

Vitari, A.C., et al. The WNK1 and WNK4 protein kinases that are mutated in Gordon’s hypertension syndrome phosphorylate and activate SPAK and OSR1 protein kinases. Biochem J 2005;391(Pt 1):17–24.

Vogel, C. and Marcotte, E.M. Insights into the regulation of protein abundance from proteomic and transcriptomic analyses. Nat Rev Genet 2012;13(4):227–232.

Voldborg, B.R., et al. Epidermal growth factor receptor (EGFR) and EGFR mutations, function and possible role in clinical trials. Ann Oncol 1997;8(12):1197–1206.

Wang, E.T., et al. Alternative isoform regulation in human tissue transcriptomes. Nature 2008;456(7221):470–476.

Woo, S., et al. Proteogenomic database construction driven from large scale RNA-seq data. J Proteome Res 2014;13(1):21–28.

Zaret, K.S. and Grompe, M. Generation and regeneration of cells of the liver and pancreas. Science 2008;322(5907):1490–1494.

Zhang, Y., et al. Prognostic implications of TOR1B expression across cancer types: a focus on basal-like breast cancer and cellular adaptations to hypoxia. J Cancer Res Clin Oncol 2024;150(6):293.

Zhou, J., et al. The function and pathogenic mechanism of filamin A. Gene 2021;784:145575.

