## Supplementary data for "PEXMap: A proteogenomic method for exon and isoform level mapping of mass spectrometry derived peptides"

Supplementary Table S1: **Detailed statistics of MS/MS peptides multi-level mapping to gene/transcript/exon/exon-exon junction against both 8-mer databases.**

| **(A) Gene-level analysis** | | |
|---|---|---|
| MS/MS peptides annotated in **PeptideAtlas** | 2,944,951 | |
| Filtered MS/MS peptides (≥8 aa, without homopolymer repeat) mapped to single genes in **PeptideAtlas** | 1,739,961 | |
| Genes mapped in **PeptideAtlas** (Genes cross-referenced in our dataset) | 15,855 (15,771) | |
| **PEXMap concordance with PeptideAtlas** | **PEXMap mapping based on** | |
| | **octamerDB** | **exonjunctionDB** |
| Total MS/MS peptides mapped by **PEXMap** | 1,723,936 (99%) | 464,033 (27%) |
| Accurately‡ mapped MS/MS peptides to single genes by **PEXMap** | 1,712,990 (98%) | 441,240 (25%) |
| Genes mapped in **PEXMap** | 15,578 (98%) | 12,226 (78%) |
| **(B) Transcript/Isoform-level analysis (peptide-to-isoform with one-to-many mapping)** | | |
| MS/MS peptides annotated to one or more transcript/isoform identifiers in **PeptideAtlas** | 2,820,860 | |
| Unique transcripts/isoforms identifiers mapped in **PeptideAtlas** (Transcripts/isoforms cross-referenced in our dataset) | 112,312 (97,443) | |
| **PEXMap concordance with PeptideAtlas** | **PEXMap mapping based on** | |
| | **octamerDB** | **exonjunctionDB** |
| MS/MS peptides mapped by **PEXMap** | 2,721,250 (94%) | 748,907 (26%) |
| Accurately‡ mapped MS/MS peptides by **PEXMap** | 2,287,817 (84%) | 592,411 (21%) |
| Unique transcripts/isoforms mapped by **PEXMap** | 85,023 (76%) | 72,592 (65%) |
| **(C) Transcript/isoform-level analysis (peptide-to-isoform with one-to-one mapping)** | | |
| MS/MS peptides annotated to **single** transcript/isoform identifier in **PeptideAtlas** | 572,037 | |
| **PEXMap concordance with PeptideAtlas** | **PEXMap mapping based on** | |

| | **octamerDB** | **exonjunctionDB** |
|---|---|---|
| Accurately‡ mapped MS/MS peptides to **single** transcript/isoform identifier by **PEXMap** | 533,486 (93%) | 140,667 (24%) |
| **(D) Exon-level analysis (exclusive to PEXMap)** | | |
| Filtered MS/MS peptides (≥8 aa, without homopolymer repeat) mapped to single genes in **PeptideAtlas** | 1,739,961 | |
| **PEXMap annotation** | **PEXMap mapping based on** | |
| | **octamerDB** | **exonjunctionDB** |
| MS/MS peptides mapped to **single exon** or **single exon-exon junction** by **PEXMap**† | 1,401,346 (81%) | 385,775 (22%) |

‡Accurately refers to identical annotations between PEXMap and PeptideAtlas.

†-MS/MS peptides accurately annotated to single genes were mapped to exons.

**Note:** All reported percentages are calculated by dividing **PEXMap** counts by the corresponding **PeptideAtlas** counts (MS peptides, genes, or transcripts).

Supplementary Table S2. **Detailed statistics of MS/MS peptides matched to liver and pancreas proteome using both octamerDB and exonjunctionDB**

| **Available MS/MS data (PeptideAtlas)** | | | | |
|---|---|---|---|---|
| **Tissues** | **Liver** | | **Pancreas** | |
| Total MS/MS peptides in **PeptideAtlas** | 3,251,966 | | 136,880 | |
| MS/MS peptides mapped to single genes in **PeptideAtlas** | 197,570 | | 80,883 | |
| **(A) Gene-level analysis** | | | | |
| **PEXMap concordance with PeptideAtlas** | **PEXMap mapping based on** | | | |
| | **octamerDB** | | **exonjunctionDB** | |
| **Tissues** | **Liver** | **Pancreas** | **Liver** | **Pancreas** |
| Mapped MS/MS peptides to single genes by **PEXMap** | 197,570 | 80,883 | 50,168 | 19,238 |
| Accurately[‡] mapped MS/MS peptides to single genes by **PEXMap** | 184,342 | 75,517 | 48,087 | 18,418 |
| **(B) Transcript/Isoform-level analysis (peptide-to-isoform with one-to-many mapping)** | | | | |
| **Tissues** | **Liver** | **Pancreas** | **Liver** | **Pancreas** |
| MS/MS peptides annotated to transcript/isoform identifiers in **PeptideAtlas** | 196,130 | 80,458 | 196,130 | 80,458 |
| **PEXMap concordance with PeptideAtlas** | **PEXMap mapping based on** | | | |
| | **octamerDB** | | **exonjunctionDB** | |
| **Tissues** | **Liver** | **Pancreas** | **Liver** | **Pancreas** |
| MS/MS peptides mapped by **PEXMap** | 191,784 | 78,762 | 49,871 | 19,142 |
| Accurately[‡] mapped MS/MS peptides | 161,824 | 67,453 | 40,167 | 15,779 |
| **(C) Transcript/isoform-level analysis (peptide-to-isoform with one-to-one mapping)** | | | | |
| **Tissues** | **Liver** | **Pancreas** | **Liver** | **Pancreas** |
| MS/MS peptides annotated to **Single** transcript/isoform identifier in **PeptideAtlas** | 47,644 | 22,672 | 47,644 | 22,672 |
| **PEXMap concordance with PeptideAtlas** | **PEXMap mapping based on** | | | |
| | **octamerDB** | | **exonjunctionDB** | |

| Tissues | Liver | Pancreas | Liver | Pancreas |
|---|---|---|---|---|
| Accurately[‡] mapped MS/MS peptides to **Single** transcript/isoform identifier to by **PEXMap** | 45,425 | 21,478 | 11,596 | 5,108 |
| **(D) Exon-level analysis (exclusive to PEXMap)** | | | | |
| **PEXMap annotation** | **PEXMap mapping based on** | | | |
| | **octamerDB** | | **exonjunctionDB** | |
| **Tissues** | **Liver** | **Pancreas** | **Liver** | **Pancreas** |
| MS/MS peptides mapped to **single exon** or **single exon-exon junction** by **PEXMap**[†] | 153,968 | 63,442 | 41,470 | 16,124 |

[‡]Accurately refers to identical annotations between PEXMap and PeptideAtlas.

[†]-MS/MS peptides accurately annotated to single genes were mapped to exons.

Supplementary Table S3. **Detailed statistics of MS/MS peptides matched to cancer proteome using both octamerDB and exonjunctionDB**

| **Category** | **Features** | **PeptideAtlas** | |
|---|---|---|---|
| **Cancer dataset composition** | Tissue samples (peptides) | 1,338,031 | |
| | Cancer cell lines (peptides) | 1,450,919 | |
| | Combined peptides (unique MS/MS peptides) | 2,104,053 | |
| **PEXMap concordance with PeptideAtlas** | | **PeptideAtlas** | **PEXMap** |
| **Gene-level mapping** | MS/MS peptides mapped to single genes | 1,152,437 | 1,117,863 |
| | Genes mapped | 14,549 | 14,296 |
| **Transcript-level mapping** | MS/MS peptides annotated to **single** transcript/isoform identifier (peptide-to-isoform with one-to-one mappings) | 326,423 | 290,516 |
| | Unique transcripts/isoforms mapped | 8,417 | 7,489 |
| **Exon-level mapping** | MS/MS peptides mapped to single exons or exon-exon junctions | - | 904,958 |

**S1. Details of exon nomenclature using ENACT framework.**

Each exon in ENACT framework is assigned unique identifier Exon Unique IDentifier (EUID), which encompass exon's attribute of relative position in gene, coding status, splice site variant, occurrence. EUID is organized into three blocks capturing coding status and contribution (Block I), inclusion frequency and position (Block II), and splice-site variation with event count (Block III). EUID is comprised of six-character code that describes exon features across three blocks.

Block I includes coding status (T: coding, U: non-coding, D: both, M: single-nucleotide coding, R: intron retention) and amino acid contribution (−2: none, −1: premature stop, 0: placeholder, 1: contributes, ≥2: multiple variants).

Block II covers inclusion frequency (G: constitutive, A: alternative, F: constitutive-like) and exon position in the gene.

Block III details splice-site variation (n: 5′, c: 3′, b: both, 0: none) and count of splice-site changes (Verma, et al., 2024).

**S2. Tissue-specific isoforms of *SCAMP3* and *ACOX1* supported by peptide mapping.**

***SCAMP3:*** *SCAMP3* (NCBI ID: 10067) gene encodes a protein involved in endosomal trafficking and membrane recycling (Fernandez-Chacon and Sudhof, 2000; Law, et al., 2012). Liver (Supplementary Fig. S1A) showed peptides (n = 18) supporting the canonical isoform (NP_005689.2) supported by exonic and their junctions (e.g., EXj 1-2 and EXj 2-3). In contrast, pancreas expressed peptides (n = 17) supported an alternative isoform (NP_443069.1), characterized by skipped exon-2 resulting in exon-exon junctions (EXj 1-3), showing tissue-specific exon usage. Quantitative proteomic studies have revealed that endocytosis-associated proteins are altered during hepatocellular carcinoma progression (Naboulsi, et al., 2016), these findings highlight that EXj peptide-based mapping enables precise identification of transcript isoforms within membrane trafficking pathways, which may be relevant in disease-associated proteomic changes.

***ACOX1***: The *ACOX1* (NCBI ID: 51) gene encodes a key enzyme in peroxisomal β-oxidation with well-established liver-enriched expression (Kumar, et al., 2024; Uhlen, et al., 2015). We observed a substantially higher number of peptides (n = 191) mapped in liver tissue (Supplementary Fig. S1B), covering multiple exons and exon-exon junctions (e.g., EXj 2–3), supporting the expression of canonical isoform (NP_009223.2). In contrast, only a small number of peptides (n = 23) were detected in pancreas, where the presence of exon-4 peptides in the absence of upstream junction suggested expression of an alternative isoform (NP_004026.2). *ACOX1* alternative splicing involving exon-3 is proposed to modulate fatty acyl substrate specificity and is prominently observed in metabolically active tissues such as liver; consistent with our observations, both isoforms are equal in length and differ by exon switching between exons 3 and 4 (Morais, et al., 2007).

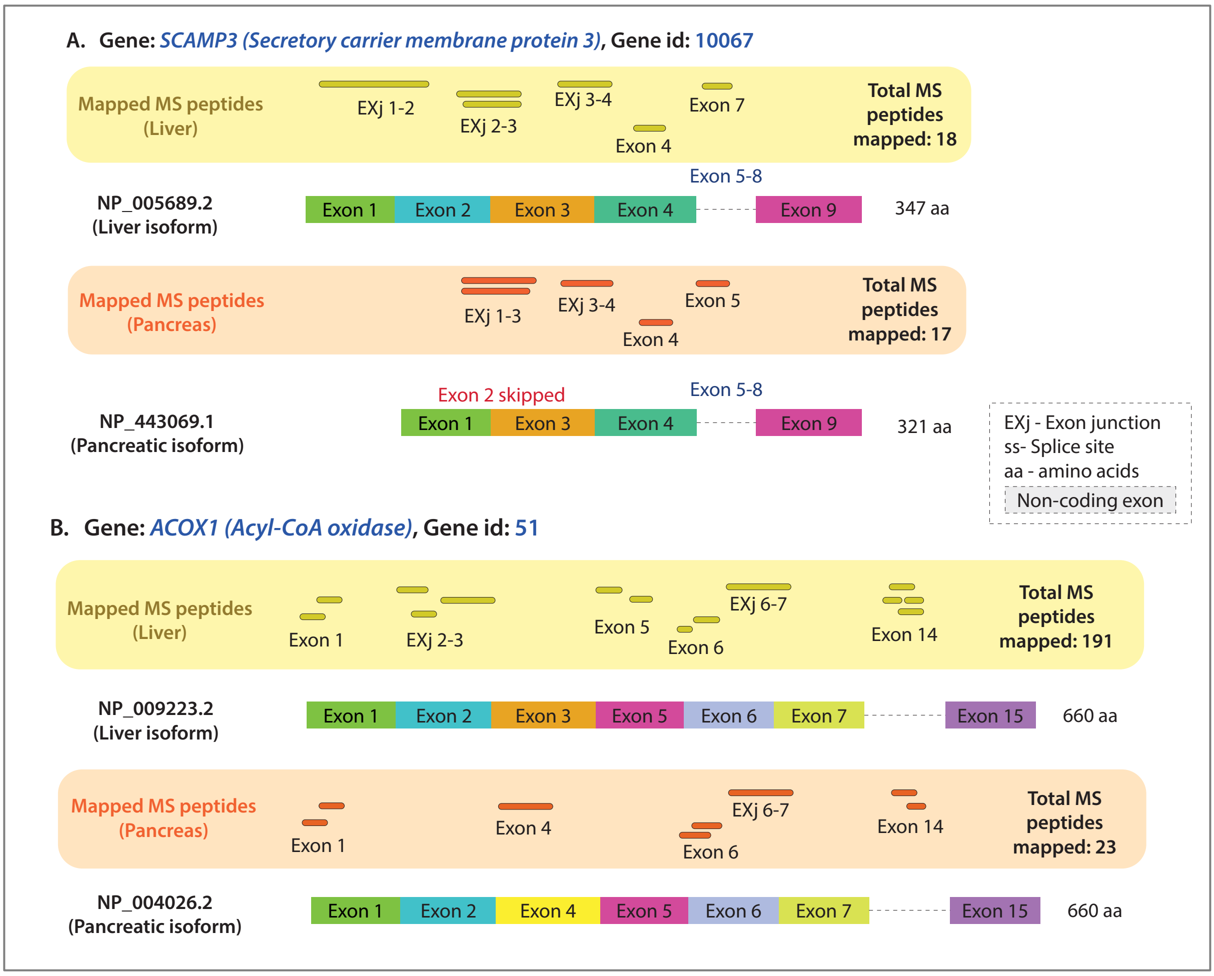


**Supplementary Figure S1: Liver- and pancreas-specific isoforms in representative genes** (A) *SCAMP3* liver isoform (NP_005689.2) includes exon 2, whereas the pancreatic isoform (NP_443069.1) shows exon-2 skipping; (B) *ACOX1* liver isoform (NP_009223.2) and pancreatic isoform (NP_004026.2) differ by differential exon usage. In each panel, annotated isoforms are shown as combinations of rectangular exon boxes, with splice-site variations indicated by altered exon boundaries. Matched MS/MS peptides are displayed as thick lines above the corresponding isoform. Numbers indicate the total mapped peptides supporting each isoform. Yellow boxes represent liver-derived peptides and orange boxes represent pancreasderived peptides. EXj denotes exon-exon junction peptides, ss indicates splice site, and dashed boxes indicate non-coding exons.

**S3. *CASP2* isoform supported by exon-level peptide evidence in cancer.**

*CASP2* (NCBI ID: 835) gene encodes *Caspase-2*, a key mediator of apoptosis that links DNA damage to cell cycle regulation and tumor suppression. Beyond its classical role, *CASP2* contributes to maintaining genomic stability and coordinating cellular stress responses, highlighting its importance in cancer biology (Kumar, 2009). In our data, MS/MS peptides mapped to exons 11–13 (Supplementary Fig. S2A), suggesting expression of an isoform (NP_116764.2) that includes these terminal exons, consistent with reports that *CASP2* splicing modulates apoptotic signaling in cancer (Shalini, et al., 2015). While it has remained unclear which *CASP2* isoform is functionally relevant in cancer, these exons are unique to a single transcript (NP_116764.2), facilitating a direct identification of transcripts at the protein level.

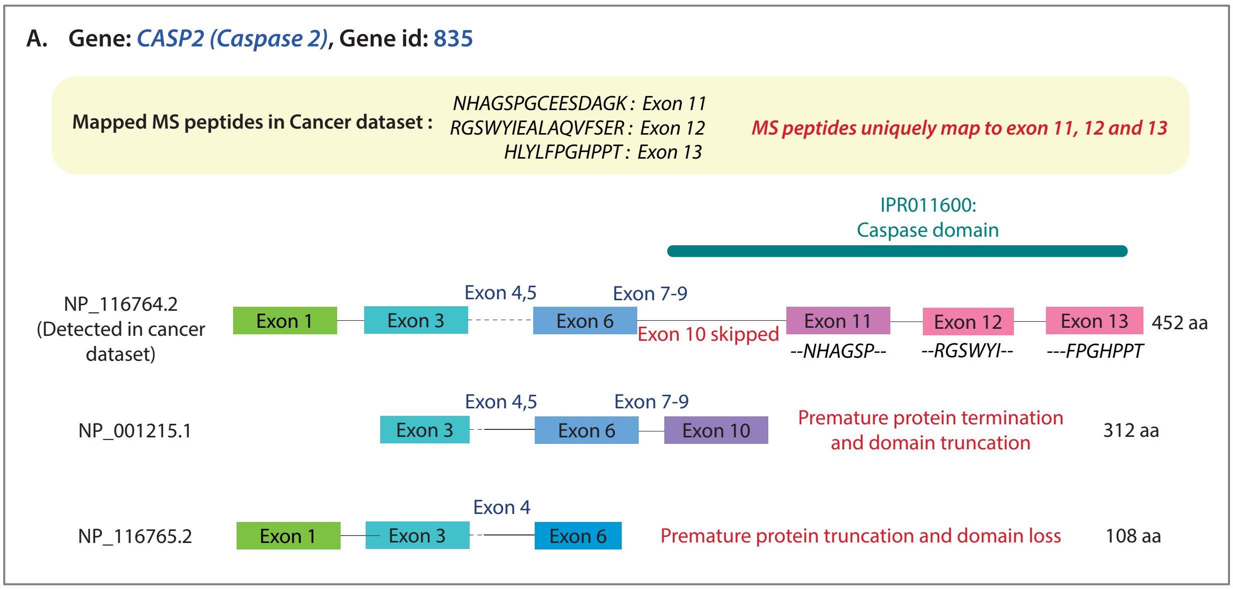


**Supplementary Figure S2: *CASP2* isoform in cancer proteome** (A) *CASP2:* peptides mapped to exons 11–13 support expression of an isoform (NP_116764.2) containing these terminal exons, consistent with alternative splicing of *CASP2* in cancer. Mapped peptide support for each isoform is indicated, and AS events are highlighted in red. Exon and peptide representations follow the schematic format used in Supplementary Fig. S1.

### References


Fernandez-Chacon, R. and Sudhof, T.C. Novel SCAMPs lacking NPF repeats: ubiquitous and synaptic vesicle-specific forms implicate SCAMPs in multiple membrane-trafficking functions. *J Neurosci* 2000;20(21):7941–7950.

Kumar, R.*, et al.* The peroxisome: an update on mysteries 3.0. *Histochem Cell Biol* 2024;161(2):99–132.

Kumar, S. Caspase 2 in apoptosis, the DNA damage response and tumour suppression: enigma no more? *Nat Rev Cancer* 2009;9(12):897–903.

Law, A.H., Chow, C.M. and Jiang, L. Secretory carrier membrane proteins. *Protoplasma* 2012;249(2):269–283.

Morais, S.*, et al.* Conserved expression of alternative splicing variants of peroxisomal acyl-CoA oxidase 1 in vertebrates and developmental and nutritional regulation in fish. *Physiol Genomics* 2007;28(3):239–252.

Naboulsi, W.*, et al.* Quantitative proteome analysis reveals the correlation between endocytosis-associated proteins and hepatocellular carcinoma dedifferentiation. *Biochim Biophys Acta* 2016;1864(11):1579–1585.

Shalini, S.*, et al.* Old, new and emerging functions of caspases. *Cell Death Differ* 2015;22(4):526–539.

Uhlen, M.*, et al.* Proteomics. Tissue-based map of the human proteome. *Science* 2015;347(6220):1260419.

Verma, P., Thakur, D. and Pandit, S.B. Exon nomenclature and classification of transcripts database (ENACTdb): a resource for analyzing alternative splicing mediated proteome diversity. *Bioinform Adv* 2024;4(1):vbae157.
